# Single-Cell Map of Dynamic Multicellular Ecosystem of Radiation-Induced Intestinal Injury

**DOI:** 10.1101/2023.03.06.531402

**Authors:** Hao Lu, Hua Yan, Yuan Xing, Yumeng Ye, Siao Jiang, Luyu Ma, Hongyan Zuo, Yanhui Hao, Chao Yu, Yang Li, Yiming Lu, Gangqiao Zhou

## Abstract

Intestine is a highly radiation-sensitive organ that could be injured during the radiotherapy for abdominal or pelvic cavity tumors. However, the dynamic change of the intestinal microenvironment related to radiation-induced intestine injury (RIII) is still unclear. Using single-cell RNA sequencing, we pictured a dynamic landscape of the intestinal microenvironment during RIII and regeneration. We showed that the multicellular ecosystem of intestine exhibited heterogeneous radiosensitivities. We revealed the distinct dynamic patterns of three subtypes of intestinal stem cells (ISCs), and the cellular trajectory analysis suggested a complex interconversion pattern among them. For the immune cells, we found that *Ly6c*^+^ monocytes can give rise to both pro-inflammatory macrophages and resident macrophages after RIII. Besides, through cellular communication analysis, we identified a positive feedback loop between the macrophages and endothelial cells, which could amplify the inflammatory response induced by radiation. Overall, our study provides a valuable single-cell map of the dynamic multicellular ecosystem during RIII and regeneration, which may facilitate the understanding of the mechanism of RIII.

## INTRODUCTION

As one of the most sensitive organs to ionizing radiation, intestine could be damaged during the radiotherapy for abdominal or pelvic cavity tumors or uncontrolled release of radioactive materials, which would lead to radiation-induced intestinal injury (RIII). Treatments of RIII are very limited and generally focused on reducing symptoms, and the curative effects are not satisfactory. RIII is largely defined as clonogenic cell death and apoptosis in the crypt cells, which results in insufficient replacement of villus epithelium, breakdown of the mucosal barrier^1, 2^, inflammation and immune abnormality^3^. Previous studies mainly focused on the roles of molecules and pathways in DNA damage, apoptosis, autophagy^4^, inflammation and immune^5^. However, the dynamics of the microenvironment during intestine injury and regeneration remains poorly understood, which limits the elucidation and treatment of RIII. Therefore, the present study aims to clarify the dynamic variation of the cellular microenvironment of RIII.

It has been widely accepted that the intestinal stem cells (ISCs) are able to replenish the whole crypt–villus axis, generating all differentiated cell types required for the physiological function of the intestine^6^. A number of ISC subpopulations in the small intestine have been identified, including Lgr5+ crypt-based columnar cells (CBCs), +4 reserve stem cells (RSCs) and revival stem cells (revSCs). Lgr5+ CBCs are considered indispensable for intestine recovery following exposure to radiation^7^. +4 RSCs, which express specific markers *Bmi1*, *Hopx* and *Tert*^8–10^, have been described as a slow dividing reserve stem cell population. Recently, a group of revSCs was identified to be extremely rare under homoeostatic conditions and arise in damaged intestines to reconstitute Lgr5+ ISCs and regenerate the intestine^11^. Irradiation causes a sharp reduction in the number of ISCs, which brings great challenges in the repair of injured intestinal epithelium. Despite the advances in our understanding of the ISCs in the past few years, a dynamic landscape of ISCs during injury and regeneration is still lacking and their interconversion relationships remain puzzling.

Immune cells also play important roles in the pathogenesis of RIII. Macrophages are crucial component of the immune system in the intestine, which modulate inflammation by secreting distinct cytokines and acting as professional phagocytes^12, 13^. Intestinal macrophages require continuous replenishment by blood monocytes and have very poor proliferative capacity, which is different from the other tissue macrophages. Exposure to radiation can significantly decreases the levels of macrophages in the damaged intestine^14^. Besides, the elucidation of the role of cross-talk between the macrophages and other cells in the initial inflammatory response and microenvironment homeostasis recovery post irradiation is also of great significance for the development of new therapeutic targets.

Recently, the single-cell RNA sequencing (scRNA-seq) has been applied to identify new cell types or cell states^15, 16^, investigate cellular plasticity and stemness of ISCs^11, 17^, or trace developmental relationships among different cell populations in the intestine^18^. However, these studies have not investigated the dynamic changes of microenvironment during intestine injury and regeneration. Here, we utilized the scRNA-seq to explore the multicellular ecosystem of homeostatic and regenerating intestine in a time-course manner. We generated transcriptomes of 22,680 single cells in the intestinal microenvironment, trying to profile the dynamics of ISCs and immune cells located in the mucous layer. Our data can be a valuable resource for further investigation on the cellular mechanism of RIII and development of potential therapy strategies.

## RESULTS

### A dynamic single-cell map of multicellular ecosystem in healthy and injured small intestine

Mice were exposed to 15 Gy of abdominal irradiation using a Co^60^ irradiator to induce intestinal injury (Figure 1A). About 20% mice died in the irradiation group within 3-6 days post irradiation (Figure 1B). The body weights of mice after irradiation exposure continued to lose in the first 7 days and then gradually increased afterward (Figure 1C). In line with this, exposed mice showed significantly reduced intestinal weight at day 3 and day 7 post irradiation as compared to unexposed mice and showed recovery at day 14. Morphological and cellular phenotype analyses also showed significant decrease of villus length, villus width, crypt depth and number of crypts at day 1 and/or day 3 post irradiation (Figures S1A–S1D). A peak of TUNEL-positive cells at day 1 post irradiation suggested that the radiation-induced cell death occurs mostly within the first 24–48 hours (Figure S1E). An increase of Ki-67-positive cells from day 3 to day 14 indicated that the day 3–14 is a key time-window for intestinal regeneration (Figure S1F).

**Figure 1.**
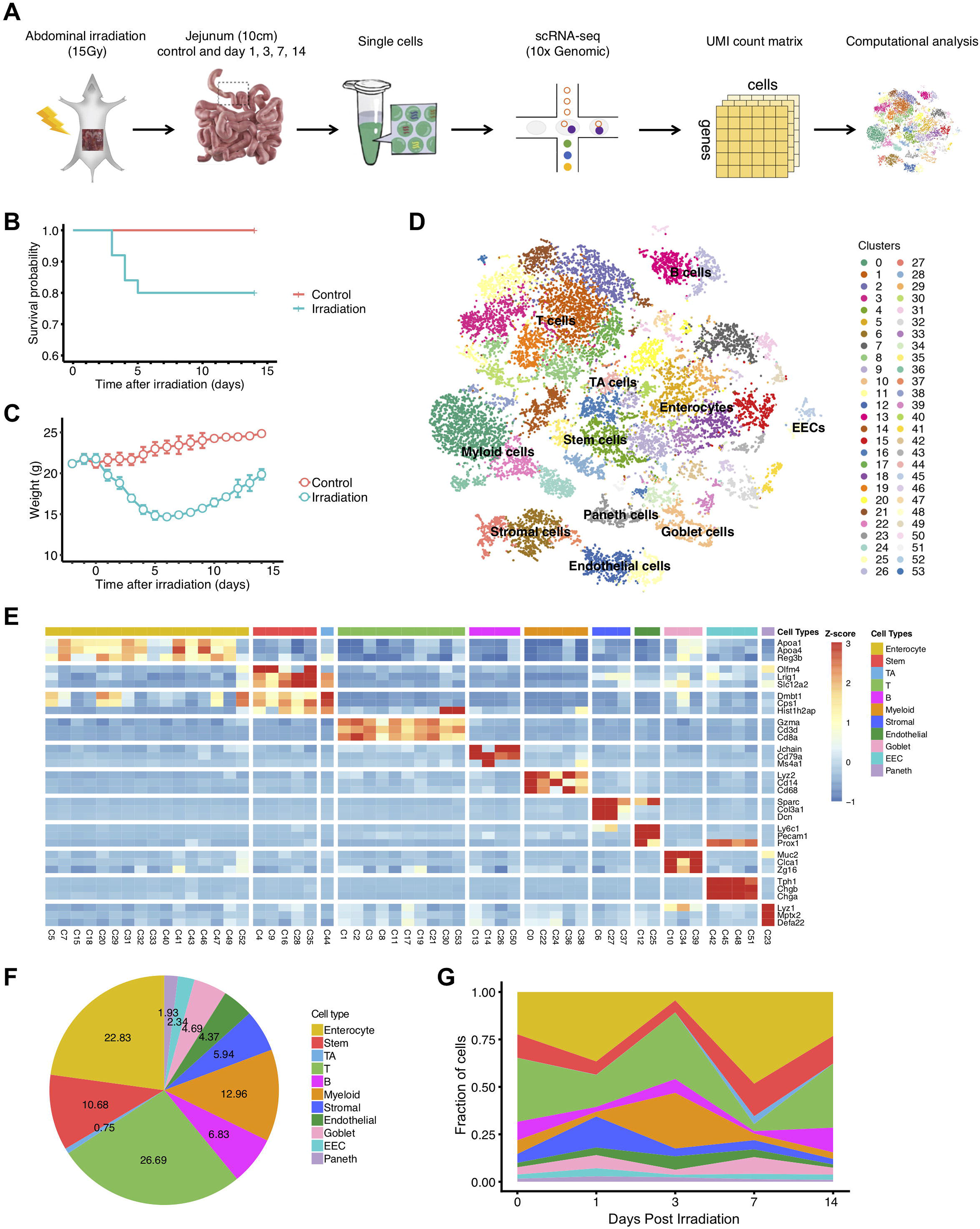
Identification of major intestinal cell types and their markers using scRNA-seq. (*A*) Overview of single-cell RNA sequencing (scRNA-seq) analysis for the irradiation-induced intestinal injury (RIII). (*B*) Survival rates of the mice exposed to 15 Gy abdominal irradiation and in control group. (*C*) Body weights of the mice exposed to 15 Gy abdominal irradiation and in control group. (*D*) tSNE projection of the 22,680 cells profiled, colored by major cell types (left), Seurat cluster (upper right) and experimental groups (lower right). (*E*) Heatmap displaying the z-score normalized mean expression of cell type-specific canonical marker genes across clusters. (*F*) Pie chart of cell type fractions in all sequenced samples. (*G*) Area chart showing the dynamic changes of the proportions of major cell types in healthy intestinal samples (Control) and exposed intestinal samples at different times after irradiation. D1, day 1; D3, day 3; D7, day 7; and D14, day 14.

To generate a dynamic single-cell map of the intestinal microenvironment related to RIII, we employed a droplet-based scRNA-seq approach to profile the transcriptomes of single cells from the intestinal tissues at day 1 (n = 4), day 3 (n = 3), day 7 (n = 3) and day 14 (n = 3) after irradiation exposure as well as the unexposed healthy intestinal tissues (n = 4) (Figure 1A). After quality filtering, we obtained the transcriptomes of a total of 22,680 single cells, with an average of 1,778 genes and 7,843 unique transcripts per cell (Figures S2A and S2B; Table S1). These transcriptomes of single cells from all samples were merged using a canonical correlation analysis (CCA)-based batch correction approach to generate a global map of cellular microenvironment of healthy and injured intestines. Shared nearest neighbor (SNN) graph-based clustering of single cells identified a total of 54 cell clusters (subtypes), which were visualized on the t-distributed stochastic neighbor embedding (t-SNE) dimensional reduction map (Figures 1D, S2C and S2D). Differentially expressed genes were calculated for each cell cluster using the Wilcoxon rank sum test.

Using canonical marker genes, we identified 10 major cell types in our dataset, including intestinal stem cells (ISCs), transit amplifying (TA) cells, enterocytes, goblet cells, enteroendocrine cells (EECs), Paneth cells, endothelial cells, T cells, B cells and myeloid cells, and most of them consist of multiple subtypes (Figures 1E and S2E), suggesting a complex cellular ecosystem of healthy and irradiation-injured intestines. We found the intestinal ecosystem changed greatly during RIII and regeneration (Figure 1F and 1G). Specifically, the stem cells and immune cells (T cells, B cells and myeloid cells) decreased markedly at day 1 post irradiation; enterocytes decreased sharply at day 3; and immune cells and endothelial cells exhibited a dramatic increase at day 3. Despite of the great alterations of intestinal microenvironments during the first 7 days post irradiation, intestinal tissues at day 14 exhibited very similar cellular composition with those non-irradiated ones. These results suggested that intestinal multicellular ecosystem disrupted by intense abdominal irradiation can be largely reconstructed within 14 days post irradiation, which is in line with our morphological observations (Figure S1).

### Intestinal multicellular ecosystem showed heterogeneity of *in vivo* radiosensitivities

The dynamic change of cellular composition in intestines before and after irradiation exposure provided an opportunity to explore the *in vivo* radiosensitivities of various cell subtypes. To quantify the radiosensitivity for each cell cluster, we compared the ratios between the observed and expected cell numbers from the intestinal samples at day 1 after irradiation. Consistent with previous studies^7, 19^, most ISC and immune cell subtypes are highly radiosensitive, exhibiting significantly lower frequencies than expected at day 1 (Figure 2A). Nevertheless, there are a few ISC and immune cell subtypes (C22, C30, C19 and C35) exhibited similar or higher frequencies than expected at day 1, suggesting they are radioresistant subpopulations. Apart from ISCs and immune cells, we found all the endothelial and stromal cell subtypes are radioresistant, while the enterocytes, goblet cells and EECs exhibit heterogeneous levels of radiosensitivity across their respective subtypes.

**Figure 2.**
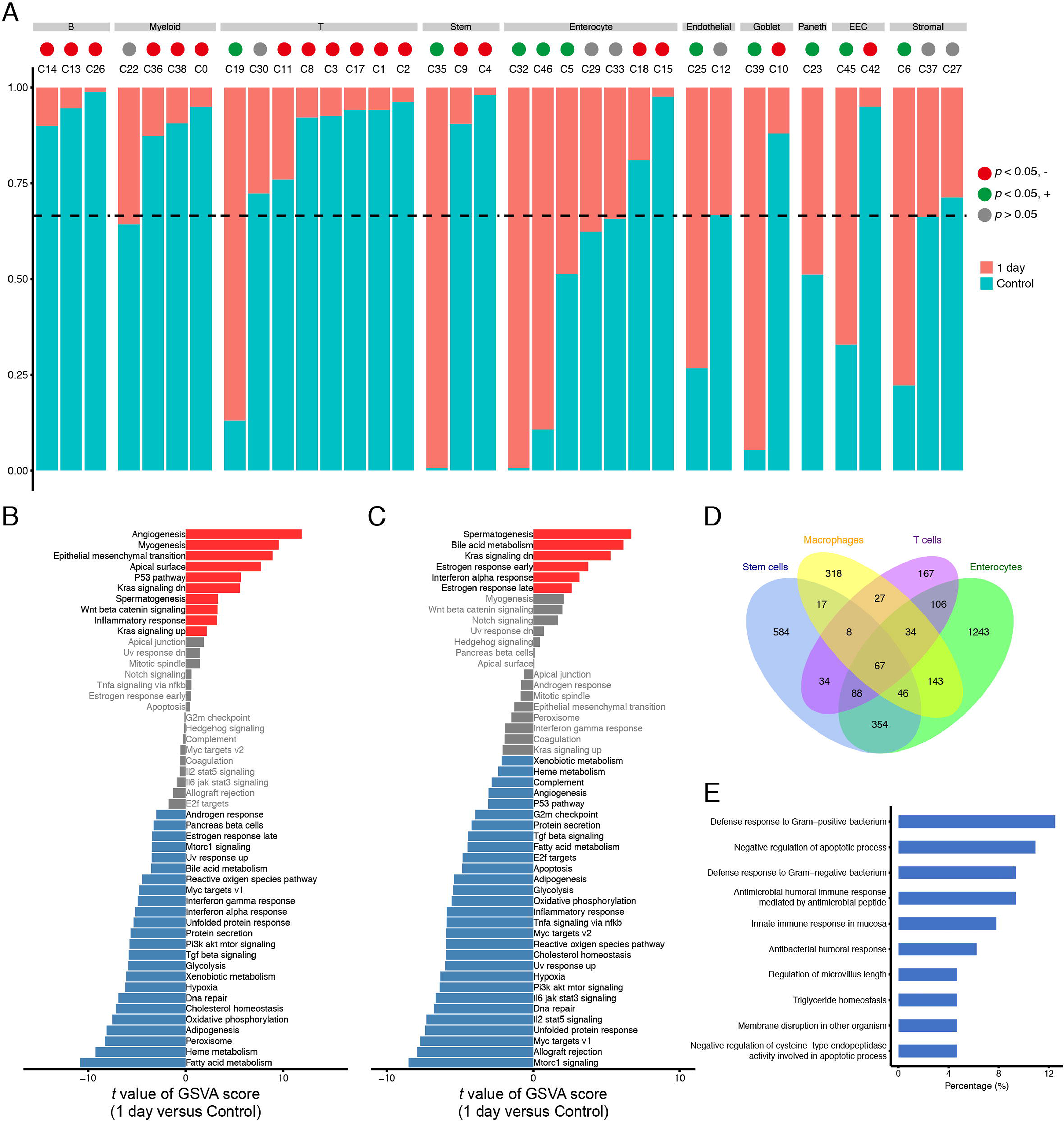
Characteristics of *in vivo* radiosensitivities of different cell types in intestine. (*A*) Bar plots showing the fraction of cells originating from the control and day 1 groups in each cluster. Only the clusters that had > 50 cells from the two groups and that showed no inter-individual difference (clusters with no more than 70% of cells from a single sample) were shown. Vertical dashed lines indicate the overall fraction of cells originating from the control group in all shown clusters. (*B*) Difference of hallmark pathway activities between stem cells from the control and day 1 (D1) groups. Shown are *t* values calculated in a linear modal comparing the pathway scores estimated by gene set variation analysis (GSVA) between cells from the two groups. (*C*) The same as (*B*) for macrophages from the control and day 1 groups. (*D*) Venn diagram showing the intersection of differentially expressed genes between cells from the control and day 1 groups for stem cells, macrophages, T cells, and enterocytes. (*E*) Top 10 significantly enriched gene ontology (GO) terms of 67 common differentially expressed genes for stem cells, macrophages, T cells, and enterocytes.

We next sought to identify the pathways activated by various cell types to confront irradiation-induced apoptosis. We focused on the top four enriched major cell types in our dataset, including the ISCs, enterocytes, myeloid cells and T cells. For each cell type, we identified pathways significantly up-/down-regulated in surviving cells at day 1 post irradiation using the whole cells from non-irradiated samples as background (Figures 2B, 2C and S3). Notably, a number of pathways, including PI3K/AKT/mTOR signaling, MYC signaling, TGF-β signaling and cell cycle-related pathways, were consistently downregulated in surviving cells of different cell types, while KRAS-down signaling pathway were consistently upregulated in surviving cells. We also investigate the genes that are upregulated in survived cells at day 1 post irradiation as compared to non-irradiated cells for the stem cells, enterocytes, myeloid cells and T cells, respectively. We identified 67 significantly upregulated genes that were shared by all the major cell types (Figures 2D and Table S2). Functional annotation showed that the upregulated genes include several genes known to be involved in defense response and negative regulation of apoptotic process (Figures 2E and Table S3).

### Dynamics of three distinct ISC subpopulations during intestinal injury and regeneration

ISCs are critical for epithelium regeneration after injury^7, 11^; however, their phenotypic heterogeneity and dynamics during intestinal injury and regeneration have not been fully characterized. We identified a total of 2,411 ISCs that were divided into five clusters (Figures 3A and 3B). Cells in clusters C4, C9, and C16 highly expressed *Lgr5*^+^ CBC signatures (*Olfm4*, *Smoc2* and *Prom1*), indicating they are *Lgr5*^+^ CBCs; additionally, cells making up these clusters showed stronger expression of ribosomal genes, suggestive of their higher transcriptional activity (Figures S4A). Stem cells in C35 expressed high levels of +4 RSC signatures (*Hopx* and *Tert*), suggesting they are a set of +4 RSCs^20, 21^. Cluster C28 was enriched for stem cells that highly expressed *Clu*, *Cxadr* and *Anxa1*, which are signatures of recently reported *Clu*^+^ revSCs^11^.

**Figure 3.**
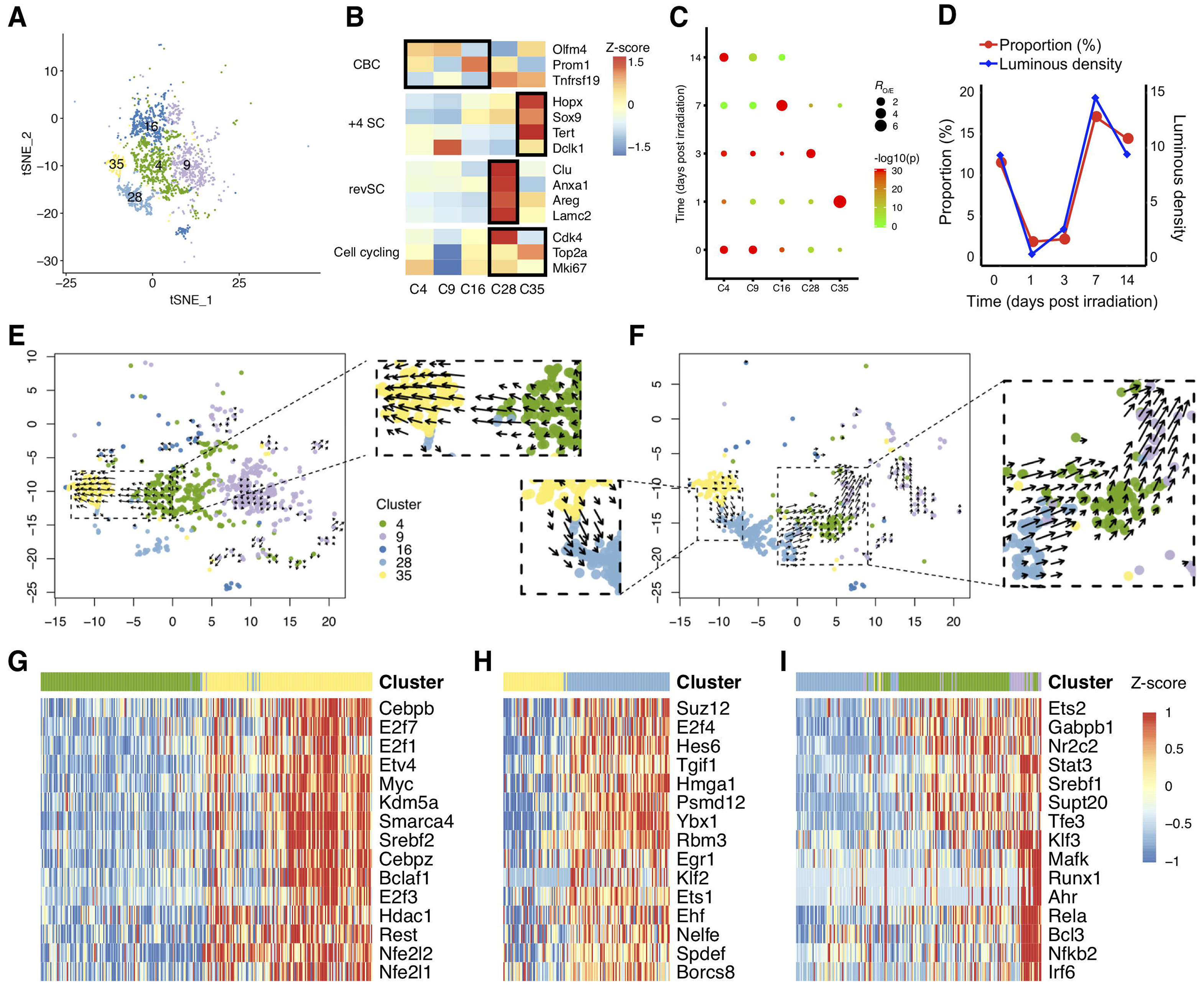
Dynamics and differentiation trajectories of intestinal stem cell (ISC) subsets in irradiation-induced intestinal injury and regeneration. (*A*) tSNE projection of 2,411 ISCs identified, colored by Seurat cluster identities. (*B*) Heatmap of the average expression of the selected ISC function-related marker genes in five ISC clusters. (*C*) Dot plots showing the ratio of observed to expected cell numbers (*R*_O/E_) of each ISC cluster in the indicated samples, with log-transformed Bonferroni-corrected *P* values in Chi-square tests. (*D*) Line charts showing the average luminous density of Olfm4 in immunostaining (blue) and the fraction of *Olfm4*^+^ CBCs (clusters 4, 9 and 16; red) in the indicated groups. (*E*) RNA velocities of ISCs from the control and day 1 groups visualized on the tSNE projection. (*F*) The same as (*E*) for ISCs from the day 1 and day 3 groups. (*G*) Heatmap depicting the estimated activity of top 15 regulons showing differential activation in ISCs along the velocity flow from the control and day 1 groups, which is depicted by a black dashed line in (*E*). Shown are normalized mean area under the curve (AUC) scores of expression regulation by each transcription factor estimated in SCENIC. Cells are ordered according to first principal component (PC1) coordinate to grasp the primary velocity orientation. (*H*, *I*) The same as (*G*) for ISCs along the velocity flow from the day 1 and day 3 groups, which is depicted by black dashed lines in the left (*H*) and right (*I*) panel in (*F*).

We then investigated the temporal dynamics of these ISC subpopulations. The proportion of *Lgr5*^+^ CBCs was drastically reduced at day 1 and day 3 and was substantially recovered at day 14 post irradiation (Figure 3C). In line with the scRNA-seq data, the integral luminous density of *in situ* hybridization with probes specific for *Olfm4* showed similar results (Figure 3D, S4B and S4C). By contrast, +4 RSCs in C35 were specifically enriched at day 1 post irradiation, suggesting they are not only radioresistant but can also be induced by irradiation (Figure 3C), in line with a previous study that detected increased lineage tracing output of +4 RSCs after irradiation exposure^22^. Distinct from *Lgr5*^+^ CBCs and +4 RSCs, *Clu*^+^ revSCs in C28 were specifically enriched at day 3 post irradiation (Figure 3C), in accordance with the study that first reported the population of *Clu*^+^ revSCs^11^. Collectively, these results showed the distinct dynamic patterns of three ISCs subpopulations. Moreover, the temporal sequential enrichment of +4 RSCs, *Clu*^+^ revSCs and *Lgr5*^+^ CBCs during this process hints at a potential sequential differentiation relationship among them.

### Differentiation trajectory analyses of intestinal stem cell subpopulations

We next sought to explore the differentiation trajectories among these heterogeneous ISC subpopulations. RNA velocity has recently emerged as a powerful approach for inferring the transition direction of a single cell to neighboring cells^23^. Here, we performed RNA velocity analysis on stem cell populations in a time-course manner. Given that only +4 RSCs were enriched at day 1, we combined them with ISCs presented under homeostatic conditions (Control-D1) or those presented at day 3 (D1-D3), respectively, to investigate the inter-subpopulation differentiation trajectories. Notably, the Control-D1 RNA velocity map showed that there is a velocity flow from *Lgr5*^+^ CBCs toward +4 RSCs (Figure 3E), in accordance with a previous study that demonstrated the interconversion between *Lgr5*^+^ CBCs and +4 RSCs^10^. This result may explain the specific enrichment of +4 RSCs at day 1 post irradiation. We observed two clear velocity flows in the D1-D3 map (Figure 3F). The first flow initiates from the +4 RSCs toward *Clu*^+^ revSCs, suggesting +4 RSCs may be able to differentiate to *Clu*^+^ revSCs. This result is not only in accordance with the specific enrichment of *Clu*^+^ revSCs at day 3 post irradiation rather than at day 1 (Figure 3B), but also consistent with the observation of a peak of CLU-tdTomato^+^ clone frequency at the +4 position from the crypt bottom^11^, where +4 RSCs are specifically enriched. The second flow is from the *Clu*^+^ revSCs toward *Lgr5*^+^ CBCs, suggesting that *Clu*^+^ revSCs could repopulate *Lgr5*^+^ CBCs when the latter were almost eliminated by irradiation, in line with the previous study^11^.

We then used SCENIC^24^ to investigate the potential key regulons along these velocity flows. We found Myc and Hdac1 were among the top transcription factors (TFs) that exhibit significantly increasing activities along the velocity flow from *Lgr5*^+^ CBCs toward +4 RSCs (Figure 3G). Myc has been reported to be required for intestinal formation and regeneration but dispensable for homeostasis of the adult intestinal epithelium^25, 26^. Here, our result suggested Myc may facilitate intestinal regeneration by promoting the conversion from *Lgr5*^+^ CBCs to +4 RSCs upon radiation-induced injury. Besides, Hdac1 was also reported to regulate intestinal stem cell homeostasis^27^. Among the top correlated TFs along the velocity flow from +4 RSCs toward *Clu*^+^ revSCs (Figure 3H), Ybx1 is a stress-activated TF and has been reported to be able to transcriptionally activate *Clu* expression by directly binds to the promoter regions of *Clu*^28^. Our data suggest Ybx1 may play important roles in the induction of *Clu*^+^ revSCs from +4 RSCs. Besides, another TF Hmga1 has been reported to amplify the Wnt signaling and expand the intestinal stem cell compartment and Paneth cell niche^29^. For the velocity flow from *Clu*^+^ revSCs toward *Lgr5*^+^ CBCs, we found Runx1 and Stat3 were among the top correlated TFs (Figure 3I). Stat3 has been shown to be indispensable for damage-induced crypt regeneration^30^ and the Runx1-Stat3 signaling pathway has been reported to regulate the differentiation of *Lgr5*^+^ CBC^31^. Taken together, our results revealed the possible differentiation trajectories among *Lgr5*^+^ CBCs, +4 RSCs and *Clu*^+^ revSCs upon intestinal injury, and identified the specific TF sets driving these differentiation paths.

### Bidirectional differentiation of peripheral monocytes into pro-inflammatory macrophages and resident macrophages

A total of 4,010 myeloid cells were identified in our dataset, which were divided into five clusters (C0, C22, C24, C36 and C38) (Figure 4A). Cells in cluster C0 expressed high levels of monocyte markers (*Cd14*, *Ccr2* and *Ly6c2*), indicating they are peripheral *Ly6c*^+^ monocytes. Cells in C22 highly expressed macrophage markers (*Cd80*, *Cd206*, *Cd64* and *F4/80*) and a set of pro-inflammatory genes (*Il1a*, *Il1b*, *Il6* and *Tnf*), suggesting they are a set of pro-inflammatory macrophages. C36 was enriched for cells expressed high levels of resident macrophage markers (*Cx3cr1* and *Fcgr1*), suggesting they are *Cx3cr1*^+^ resident macrophages. Besides, we also identified a cluster of neutrophils (C24) highly expressing neutrophil markers (*Ly6g*, *Retnlg* and *S100a9*) (Figure 4B).

**Figure 4.**
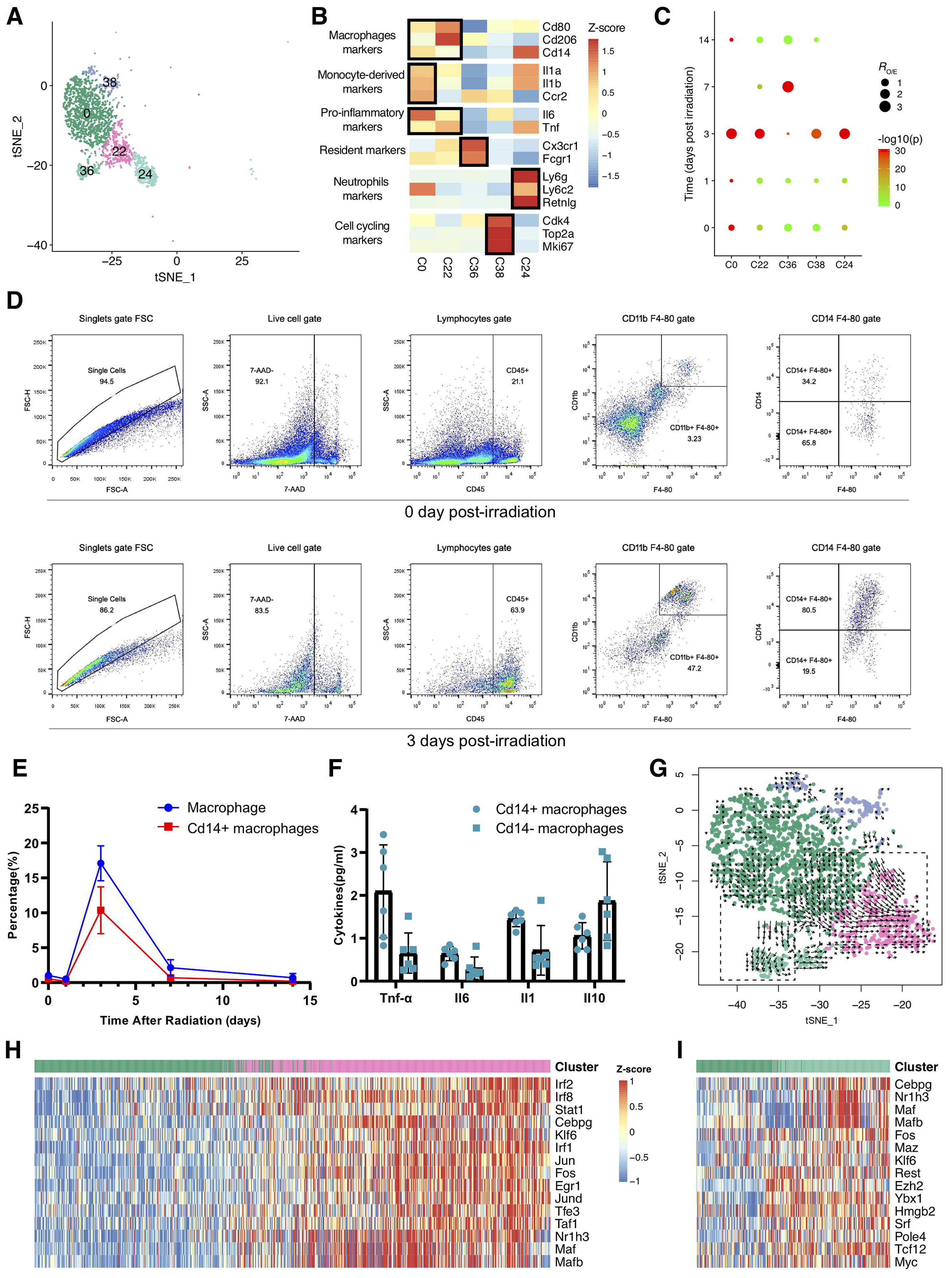
The characteristics and roles of macrophage subsets in the irradiation-induced intestinal inflammation. (*A*) tSNE projection of 2,926 myeloid cells identified, colored by Seurat cluster identities. (*B*) Heatmap of the average expression of the selected myeloid cell function-related marker genes in five myeloid cell clusters. (*C*) Dot plots showing the ratio of observed to expected cell numbers of each myeloid cell cluster in the indicated samples, with Bonferroni-corrected P values by Chi-square tests. (*D*) Flow cytometry analyses showing the percentages of Cd14^+^ inflammatory macrophages at control and 3 days post irradiation groups. (*E*) Dynamic change of the percentages of Cd14+ macrophages at control, 1, 3, 7 and 14 days post irradiation groups. (*F*) Expression levels of inflammatory cytokines by Cd14^+^ and Cd14^-^ macrophages in intestine 3 day post irradiation. (*G*) RNA velocities of four macrophage clusters visualized on the tSNE projection. (*H*, *I*) Heatmap depicting the estimated activities of top 15 differentially activated regulons along the velocity flow, which was depicted by black dashed lines in the right (*H*) and left (*I*) panel in (*G*). Shown are normalized mean area under the curve (AUC) scores of expression regulation of each transcription factor estimated by SCENIC. Cells are ordered according to first principal component (PC1) coordinate to grasp the primary velocity orientation.

We then investigated the temporal distribution of these myeloid subpopulations. We found that *Ly6c*^+^ monocytes (C0), pro-inflammatory macrophages (C22) and the neutrophils (C24) were highly enriched at day 3 post irradiation (Figure 4C). The temporal distributions of pro-inflammatory macrophages after irradiation were confirmed by flow cytometry, and their expression of pro-inflammatory cytokines were measured (Figures 4D-4F). In contrast, *Cx3cr1*^+^ resident macrophages in C36 showed depletion at day 1 and day 3 post irradiation and became abundant at day 7, suggesting resident macrophages are radiosensitive and could rapidly recover after irradiation.

Using RNA velocity analysis, we found that there are two differentiation paths initiated from *Ly6c*^+^ monocytes towards pro-inflammatory macrophages and *Cx3cr1*^+^ resident macrophages, respectively (Figure 4G). Although the alternative outcomes of *Ly6c*^+^ monocytes to be resident and pro-inflammatory macrophages has been previously reported in inflammatory bowel disease (IBD)^32^, our result shows that this bidirectional differentiation pattern may be import for RIII recovery. Further, SCENIC analysis was performed to investigate the potential key regulons driving these two differentiation paths, respectively. We found these two differentiation paths shared several top correlated TFs, including Cebp, Maf, Mafb and Nr1h3 (Figure 4H and 4I). CCAAT/enhancer-binding protein (Cebp), Maf and Mafb transcription factors has been reported to be important for monocyte-to-macrophage differentiation^33^. Nuclear receptor Nr1h3 (LXRα) has also been reported to be a major regulator of macrophage development^34^. Besides the shared TFs, the two differentiation paths were also driven by specific factors. Activities of Jun, Stat1, Irf1 and Irf2 were specifically upregulated along the velocity flow from *Ly6c*^+^ monocytes to pro-inflammatory macrophages (Figure 4H), while Myc and Myc-associated zinc finger (Maz) were specifically upregulated along the velocity flow from *Ly6c*^+^ monocytes to resident macrophages (Figure 4I), in accordance with the key role of Myc in the self-renewal activity of macrophages^35^. Together, these results suggested that peripheral monocytes could differentiate into both pro-inflammatory macrophages and resident macrophages after RIII, and these two differentiation paths may be driven by different key TFs.

### A positive feedback loop between the macrophages and endothelial cells amplifies inflammatory response upon RIII

To assess the roles of various cell types in the inflammatory response induced by irradiation, we then investigated the cell-cell communications (CCC) in the intestine microenvironment. We found that the overall and the myeloid cell-derived interaction strength showed a sharp decrease in day 1 post irradiation and a remarkable increase in day 3 post irradiation (Figure 5A). Given the significant enrichment of myeloid cells in day 3, we then compared the CCC between day 3 and day 1 in more details. Results revealed that the interaction number and strength among myeloid, endothelial and stromal cells exhibited strong upregulation in day 3 compared to day1 (Figure 5B). It is, moreover, noteworthy that myeloid cells showed significantly increased cellular interaction in signaling pathways relevant to leukocyte migration and adhesion, such as ICAM, ITGAL-ITGB2, SELPLG and SELL signaling (Figure 5C and S5). It is reasonable to postulate that the high-volume infiltration of myeloid cells in day 3 might be induced by endothelial and stromal cells.

**Figure 5.**
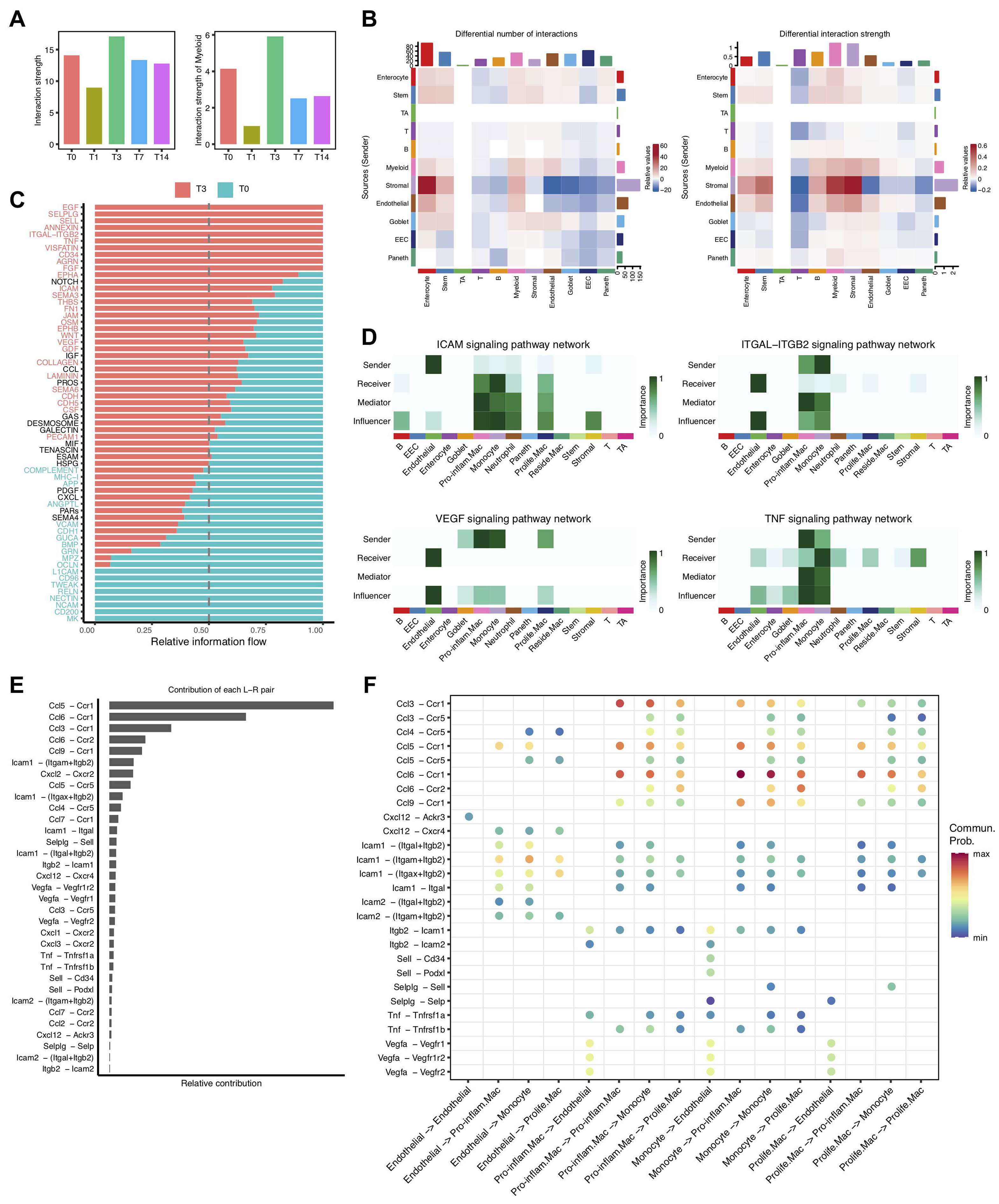
Cellular interaction between the myeloid and endothelial cells. (*A*) The overall (lift) and myeloid cell-derived (right) interaction strength at different times post irradiation. (*B*) Heatmap depicting the differential interaction numbers (left) and strength (right) among major cell types in day 3 group compared to those in control group. (*C*) Bar graph showing the overall information flow of each signaling pathway in day 3 and control group. (*D*) Heatmap depicting the network centrality scores for each cell type in the selected signaling pathways. (*E*) Bar graph showing the contribution of each ligand-receptor pair to the myeloid cell migration and activation relevant signaling pathways. (*F*) Dot plot showing the significant ligand-receptor pairs associated with the myeloid cell migration and activation relevant signaling pathways.

We then examined the CCC associated with different myeloid cell populations. We found that ICAM and ITGAL-ITGB2 signaling pathways exhibited remarkable enrichment between endothelial cells and monocytes and pro-inflammatory macrophages (Figure 5D). Investigation of specific ligands and receptors revealed that endothelial cells expressed high levels of adhesion molecules such as *Icam1*and *Icam2*, to recruit monocytes and pro-inflammatory macrophages(Figure 5E and 5F), in accordance with previous studies^36^. Besides the adhesion molecules, we found these endothelial cells also expressed high levels of chemokine *Ccl5* and *Cxcl12* to recruit pro-inflammatory macrophages (Figure 5E and 5F). Interestingly, the pro-inflammatory macrophages expressed high levels of vascular endothelial growth factor (*Vegf*), which could, in turn, enhance the endothelial cell survival and proliferation^37^ (Figure 5E and 5F). The interactions between the activated endothelial cells and pro-inflammatory macrophages could amplify the proliferation of both populations, which is consistent with the specific enrichment of pro-inflammatory macrophages and endothelial cells at day 3 post irradiation. We speculate the amplification effect is critical for the quick induction of inflammation at early stage of intestinal injury.

We also sought to identify the pathways that are upregulated upon endothelial cells activation during RIII. Compared to endothelial cells in homeostatic intestine, the activated endothelial cells at day 3 post irradiation were shown to be upregulated in inflammatory response, TNF-α signaling via NF-κB, EMT and KRAS signaling pathways (Figures S6). NF-κB has been reported to play a pivotal role in the inducible expression of cytokines in inflammatory response induced by irradiation^38^. Besides, we found KRAS signaling pathway may also play an important role in inflammatory cytokines induction. In addition, the upregulation of EMT pathway indicates that irradiation could induce the epithelial-to-mesenchymal transition of endothelial cells, which may be involved in intestinal fibrosis after radiation exposure^39^. Collectively, these results showed that the cellular interactions between macrophages and endothelial cells could achieve a quick amplification of inflammation upon the RIII.

## DISCUSSION

The dynamics of cellular microenvironment during the radiation-induced intestinal injury and regeneration remains largely uncharacterized. In this study, by combining the single-cell transcriptome profiling with temporal distribution analysis, lineage reconstruction, TF profiling and cellular interaction analyses, we provided a comprehensive dynamic landscape of the cellular microenvironment during intestinal injury and regeneration.

The radiosensitivity of cells varies greatly, mainly depending on the degree of cell differentiation, cell proliferation ability, metabolic status and surrounding environment^3, 40^. Currently, it is widely accepted that the lymphocytes, hematopoietic cells, small intestinal crypt cells and germ cells are highly radiosensitive cell types, but these kinds of cells are composed with multiple subtypes with different biological characteristics. In this study, based on scRNA-seq, we assessed the *in vivo* radiosensitivity of different cell subtypes in the intestinal microenvironment. Our data not only showed that diverse cell subtypes in the intestinal microenvironment exhibited highly heterogeneous levels of radiosensitivity, but also revealed cross-cell consistency of pathway activation for various cell types to confront the radiation-induced apoptosis.

Much attention has been focused on ISC injury and regeneration post irradiation^7, 11, 22, 41^. It is generally accepted that Lgr5+ stem cells were sensitive to irradiation injury while +4 stem cells show relatively lower sensitivity^8, 10, 22^, we here reported a comprehensive survey on the phenotype and dynamics of diverse ISC subsets in different phases after high-dose irradiation. Generally, we identified three subpopulations of ISCs, including *Lgr5*^+^ CBCs, +4 ISCs as well as a cluster of *Clu*^+^ revISCs reported recently^11^, with distinct dynamic characteristics. Additionally, our data provides new insights into the interconversion relationships among the three ISC subpopulations. Although a previous study has shown *Lgr5*^+^ CBCs could give rise to +4 RSCs in culture^10^, our data suggests this conversion could be induced *in vivo* by intestinal injury. Besides, we also showed that *Clu*^+^ revISCs may originate from *Lgr5*^+^ CBCs and +4 revISCs, although further studies are still needed to test and verify the transformation processes.

Exposure to irradiation could also strongly affect immune system responses in the intestine, which were essential for maintaining mucosal homeostasis. In this study, we observed that *Cx3cr1*^+^ resident macrophages are drastically reduced at 1 day post irradiation and were then followed by a massive influx of monocytes and macrophages into the injured intestine. Our data suggested that the bidirectional differentiation of peripheral *Ly6c*^+^ monocytes to pro-inflammatory macrophages or *Cx3cr1*^+^ resident macrophages may be important for balancing the radiation-induced inflammation and resident macrophages recovery. We also showed that different TFs may contribute to the alternative fates of *Ly6c*^+^ monocytes, which could be potential intervention targets for modulating the early stage inflammation induced by radiation.

In addition, we found the cellular interacting patterns varied greatly at different times during RIII and recovery; especially, the massive interactions between the macrophages and endothelial cells at early inflammation stage aroused our attention. Our results showed that the cross-talk between pro-inflammatory macrophages and activated endothelial cells could achieve a quick amplification of inflammation upon RIII. This finding suggested that intervention in the interaction between these two cells might alleviate early inflammation.

In summary, based on scRNA-seq, our research refined the radiosensitivity of small intestinal cells in the cell subtype level. Besides, our dataset revealed the dynamic patterns of three ISC subpopulations and their interconversion relationships. Additionally, we showed the bidirectional differentiation of peripheral monocytes into pro-inflammatory macrophages and resident macrophages, and the amplifying communicating relationships between macrophages and endothelial cells during the inflammatory response upon RIII. These findings may provide new views on the cellular and molecular mechanisms of RIII.

## MATERIALS AND METHODS

### Ethics statement

The study was approved by the institutional review board of Beijing Institute of Radiation Medicine (protocol ID: IACUC-DWZX-2020-623). The work submitted in this article was solely completed by the teams of Gangqiao Zhou, Yiming Lu and Yang Li, and it is original. Excerpts from others’ work have been clearly identified and acknowledged within the text and listed in the list of references.

### Mice and groups

Wild-type male C57BL/6J mice (weight of 20 ± 2g) were purchased from Beijing Vital River Laboratory Animal Technology Co., Ltd. (Beijing, China). All the mice were bred in a specific pathogen-free environment under conditions of constant temperature of 22 ± 1°C, relative humidity of 60%, and regular dark-light schedule (lights on from 7 a.m. to 7 p.m.) at the Experimental Animal Center of the Beijing Institute of Radiation Medicine, China. A total of 97 mice were used in this study, including 78 random mice receiving abdominal irradiation and 19 random mice as negative controls. The irradiation exposed mice were then randomly divided into four groups and intestinal samples were collected and used for experiments at day 1, 3, 7 and 14 post irradiation, respectively. Specifically, 17 mice were used for scRNA-seq (n = 3 or 4 for each group), 20 mice for flow cytometry (n = 3 for each group, with additional 5 mice for flow cytometry cell sorting at day 3), 30 mice for immunohistochemistry (n = 6 for each group), and 30 mice for *in situ* hybridization analyses (n = 6 for each group).

### Abdominal irradiation of mice

Mice were anaesthetized with an intraperitoneal injection of 0.5% pentobarbital (43 mg/kg body weight) and exposed to a single dose of 15 gray (Gy) abdominal irradiation (from the xiphoid process to the pubic symphysis with lead bricks to cover the other parts of the mice) using a Co^60^ irradiator to induce intestinal injury *in vivo*. All experiments were repeated at least twice with n = 6 mice in each group, except for the scRNA-seq, in which each group contain n = 3 or 4 mice. Mice were monitored for up to 15 days, and the changes in small intestine lengths and weights were recorded on day 0, 1, 3, 7, 14 post irradiation.

### Morphological analyses of mice villus and crypt

Mice were monitored for up to 15 days. The mice in control and irradiation groups were sacrificed on day 1, 3, 7, 14 following irradiation, and the intestine tissues were harvested and fixed in a 10 % neutral buffered formalin solution. The fixed samples were dehydrated, cleared and permeated with paraffin in a tissue processor, and subsequently embedded in paraffin blocks using an embedding system. Paraffin-embedded samples were sectioned at a 5-mm thickness and stained with hematoxylin & eosin (HE) staining. The slides were imaged at 50 ×, 100 × and 400 × magnification, respectively, using an Olympus BX51 microscope (Japan). The villus length and width, crypt depth, thickness of muscular layer and number of crypts per intestinal length were measured with ImageJ (NIH). The villus length was measured from the top to the base of the villus at the entrance to the intestinal crypt. The villus width was measured at half of its length. The crypt depth was measured from the depth of the invagination to the adjacent villi. For each group, at least 30 well-oriented villi were measured and the mean value was calculated.

### Single-cell sequencing library construction

To isolate single cells, duodenums were amputated (about 5 cm beyond the pylorus) and 10 cm jejunum segments following the incision of the mice on day 0, 1, 3, 7, 14 after irradiation were washed in cold PBS, cut longitudinally into roughly 2-mm-long pieces and were isolated using the Liver Dissociation Kit (mouse) and GentleMACS Dissociator (Miltenyi Biotec, Germany) according to the manufacturer’s instructions. Tissues were filtered through a 40-μm-mesh cell strainer on ice, pelleted by centrifugation at 4 °C and washed twice with the ice-cold regular medium to remove the debris. An estimated 5,000 single cells per sample were then subjected to 10x Genomics single-cell isolation and RNA sequencing following the manufacturer’s recommendations. Illumina HiSeq 3000 was used for deep sequencing. Two technical replicates were generated per sorted cell suspension.

### Analysis of scRNA-seq data

The Cell Ranger software (version 2.2.0) provided by 10x Genomics was used to align the reads from droplet-based scRNA-seq to an indexed mouse genome (mm10, NCBI Build 38), generating a digital gene expression matrix (UMI counts per gene per cell) for each sample. Expression matrices for all samples were filtered, normalized, integrated and clustered using the standard Seurat (version 2.3.4) package procedures. More exactly, for the first quality-control (QC) step, we removed the genes detected in less than three cells and cells with below 500 or over 8,000 expressed genes and over 10% UMIs derived from mitochondrial genome. After applying these QC criteria, a total of 22,680 cells and 19,588 genes in total remained and were included in the following analyses. Then the expression matrices were log-normalized and scaled to remove the unwanted variation from the total cellular read count and mitochondrial read count, as implemented in Seurat’s *NormalizeData* and *ScaleData* functions. To integrate datasets from different samples, we used a subset of highly variably expressed genes to perform the canonical correlation analysis (CCA). First, the top 1,000 genes with the largest dispersion in each dataset were selected; then the genes from all 17 datasets were intersected to determine an overlap-gene set. We used the overlap-gene set as variable genes to implement CCA through Seurat’s *RunMultiCCA* function, which returned an integrated Seurat objects with canonical correlation vectors. Seurat’s *AlignSubspace* function was employed to align the top ten dimensions in CCA subspaces and generate the *cca.aligned* dimensional reduction. With the first ten components of the dimensional reduction, we then performed a shared nearest neighbor (SNN) modularity optimization clustering method with the Louvain algorithm as implemented in the *FindClusters* function, which finally identified 54 clusters of different cell types or subtypes, and different clusters of cells were visualized using a further t-distributed Stochastic Neighbor Embedding (tSNE) dimensionality reduction.

### Immunohistochemistry assay

The 5 μm thick sections from the paraffin-embedded small intestine sections were deparaffinized and rehydrated using xylene and ethanol and boiled for 15 minutes (min) in 10 mM citrate buffer solution (pH 6.0) for antigen retrieval. The sections were then immersed in a 3% hydrogen peroxide solution for 10 min to block the endogenous peroxidase. Slides were incubated with goat serum for 10 min and then with the primary antibody anti-Ki67 (#ab16667, 1:200; Abcam). A horse radish peroxidase (HRP)-based signal amplification system was then hybridized to a goat anti-rabbit IgG (H&L) secondary antibody (#PV9001; ZSBIO) followed by colorimetric development with diaminobenzidine (DAB). Positive cells were counted in the crypt and villi at 30 randomly selected position per group with ImageJ.

### TUNEL assay

Apoptotic cells were identified by terminal deoxynucleotidyl transferase-mediated dUTP nick end-labeling (TUNEL) staining using the *In Situ* Cell Death Detection Kit (Roche) according to the manufacturer’s protocol. Briefly, the paraffin-embedded sections were prepared the same as hematoxylin & eosin (HE) staining. After drying and deparaffinized, the section was treated by proteinase K (20 μg/mL; Roche, Swiss) for l0 min to make the cell membrane permeable, and then treated by the mixed reaction solution for TUNEL reaction. After treatment with biotin-labeled HRP, DAB chromogen was used to render the color. TUNEL-positive cells were identified, and their numbers were counted within a defined area (μ2) using an Olympus BX51 microscope (Japan).

### *In situ* hybridization

*In situ* hybridization (ISH) assays for *Olfm4* were performed with the RNAscope kit (Advanced Cell Diagnostics, California, USA) according to the manufacturer’s instructions. Five μm formalin-fixed, paraffin embedded tissue sections or 8 μm OCT frozen were pretreated with heat and protease digestion prior to hybridization with target probes (Advanced Cell Diagnostics). An HRP-based signal amplification system was then hybridized to the target probes followed by colorimetric development with DAB or Fitc, and Cy3. The housekeeping gene *ubiquitin C* (*UBC*) was served as a positive control and the *dapB* gene, which is derived from a bacterial gene sequence, was used as a negative control.

### Identification of marker genes

To identify the marker genes for each of the 54 clusters of different cell types or subtypes, we contrasted the cells from each cluster to cells from all the other clusters using the Seurat’s *FindMarkers* function. The marker genes were required to: (1) have an averaged expression in the current cluster that is at least 0.25-fold (log-scale) larger than that in all other clusters; and (2) show expression in at least 10% of cells in either of the two comparative populations.

### Flow cytometry

Single cells prepared as above were resuspended in phosphate buffered saline (PBS). For macrophages, the antibodies used were against Cd45, F4/80, Cd11b and Cd14, and cells were stained with 7-AAD to differentiate live cells. Antibody staining was performed at RT for 30 min before washing cells twice with PBS. The 7-AAD was added to the final FACS medium for 10 min before flow cytometry analyses and/or isolation of required single-cell populations. These cells were gated on live cells (those that were 7-AAD negative) and CD45 positivity. Subsequent determinations included the presence of F4/80 and Cd11b followed by the presence or absence of Cd14. The cells were analyzed on a BD FACSCanto II flow cytometer (BD, New Jersey, USA) utilizing FACS Diva Software (BD).

### RNA velocity analyses

RNA velocity of single cells was estimated as previously reported^23^. More exactly, the spliced and unspliced transcript reads were calculated based on the CellRanger output using the velocyto command line tool with run10x subcommand. Transcript counts from different samples were merged and genes with an average expression magnitude < 0.5 (for spliced transcripts) or < 0.05 (for unspliced transcripts) in at least one of the clusters of each cell type were then removed. Cell-to-cell distance was calculated using Euclidean distance based on the correlation matrix of the *cca.aligned* dimensional reduction from Seurat. RNA velocity was estimated using gene-relative models, with k-nearest neighbor cells pooling of 20 and fit quantile of 0.02. Velocity fields were then visualized in the tSNE dimensionality reduction from Seurat.

### Gene set variation analysis (GSVA)

To assess the relative pathway activity on the level of individual cells, we conducted the gene set variation analysis (GSVA) with the GSVA package (version 1.32.0)^42^. The 50 hallmark gene sets (version 6.2) representing specific well-defined biological states or processes were obtained from the Molecular Signatures Database (MSigDB)^43^, and genes in each set were converted to orthologous genes in mouse with g:Profiler^44^. GSVA was performed in the *gsva* function with standard settings. We then fit a linear model to the output gene set-by-cell pathway enrichment matrix, implemented in the *lmFit* function of limma package (version 3.40.6), to detect the differentially enriched pathways.

### SCENIC analysis

The pySCENIC (version 0.9.11)^45^ analysis was run as described for the stem cells and myeloid cells. The normalized expression matrix exported from Seurat was set as the input of pySCENIC pipeline. In addition, the genes with less than 200 UMI counts or detected in less than 1% of the cells in the corresponding cell types were removed as noise. Two transcription factor ranking databases, namely the TSS+/-10kb and *500bpUp100Dw* databases, were used for inference of co-expression modules and identification of direct targets. A regulon-by-cell matrix was then calculated to measure the enrichment of each regulon as the area under the recovery curve (AUC) of genes defining the regulon. To identify the key regulons driving the differentiation trajectories among the stem or myeloid cell subpopupation, cells along each RNA velocity flow were selected and their tSNE coordinates were converted using principal components analysis (PCA) to capture the differentiation process on the first principal component (PC1). Differential enrichment was calculated on the SCENIC AUC matrix along the PC1 coordinates for each trajectory with a general additive model implemented in the gam package (version 1.16.1).

### Cell-cell communication analysis

We used CellChat (Version 1.6.1)^46^ to infer the cell-cell communication among different cell types following the tutorials in the CellChat github repository. In brief, a CellChat object was constructed based on the normalized expression profile at each time point post irradiation. Cellular communication probability was then computed with the default parameters and cell-cell communication with < 10 cells in the corresponding cell groups was filtered out. Communication probability on signaling pathway level was further inferred by summarizing the communication probabilities of all ligands-receptors interactions associated with each signaling pathway.

### Statistical analyses

GraphPad Prism 5 and R (version 3.6.0) were used to perform the statistical analyses. The Kaplan-Meier method was used to analyze the animal survival curves. The Chi-square test was applied to analysis the distribution of cells at different times before and after irradiation. *P*-values of multiple testing were corrected by Bonferroni correction method and an adjusted *P*-value < 0.05 was considered to be statistically significant.

### Data availability

The scRNA-seq dataset generated in this study are available at the National Center of Biotechnology Information’s Gene Expression Omniobus database under the following accession number: GSE165318.

## Acknowledgements

This work was funded by the General Program (32270714, 81573251 and 81672369) of the Natural Science Foundation of China (www.nsfc.gov.cn), Beijing Nova Program (20180059), National Key R&D Program of China (No. 2017YFA0504301), Chinese Key Project for Infectious Diseases (No. 2018ZX10732202 and 2017ZX10203205) and Beijing Institute of Radiation Medicine (BIRM) Innovation Fund (BIOX0201).

## Authors contributions

G.Z., Y.Lu. and Y.Li was the principal investigators who conceived and designed the study, obtained financial supports and approved the final version of the manuscript; H.L., H.Y., S.J. and L.M. performed the data analyses; H.Y. conducted most of the cell sorting and functional experiments; H.Y., Y.X., Y.Y., H.Z., Y.H. and C.Y. performed abdominal irradiation of mice and collected intestinal samples; G.Z., Y.Lu. and Y.Li drafted the manuscript. All the authors read and approved the final version of the manuscript.

## Competing interests

The authors disclose no conflicts.

## Notes

### Competing Interest Statement

The authors have declared no competing interest.

### Summary of Updates

We have performed cell-cell communication analysis with a new pipeline (namely CellChat) to illustrate the intercellular interactions and rephrased the 6th subsection in the result section.

## REFERENCES

1. Shukla PK, Gangwar R, Manda B, Meena AS, Yadav N, Szabo E, et al. Rapid disruption of intestinal epithelial tight junction and barrier dysfunction by ionizing radiation in mouse colon in vivo: protection by N-acetyl-l-cysteine. American Journal of Physiology-Gastrointestinal and Liver Physiology. 2016;310:G705–G715.

2. Hauer-Jensen M, Wang J, Denham JW. Bowel injury: current and evolving management strategies, In Seminars in Radiation Oncology, Elsevier, 2003.

3. Hauer-Jensen M, Denham JW, Andreyev HJ. Radiation enteropathy--pathogenesis, treatment and prevention. Nat Rev Gastroenterol Hepatol. 2014;11:470–9.

4. Qu W, Zhang L, Ao J. Radiotherapy induces intestinal barrier dysfunction by inhibiting autophagy. ACS Omega. 2020;5:12955–12963.

5. Zhao Z, Cheng W, Qu W, Shao G, Liu S. Antibiotic alleviates radiation-induced intestinal injury by remodeling microbiota, reducing inflammation, and inhibiting fibrosis. ACS Omega. 2020;5:2967–2977.

6. Spit M, Koo BK, Maurice MM. Tales from the crypt: intestinal niche signals in tissue renewal, plasticity and cancer. Open Biol. 2018;8.

7. Metcalfe C, Kljavin NM, Ybarra R, de Sauvage FJ. Lgr5+ stem cells are indispensable for radiation-induced intestinal regeneration. Cell Stem Cell. 2014;14:149–59.

8. Montgomery RK, Carlone DL, Richmond CA, Farilla L, Kranendonk ME, Henderson DE, et al. Mouse telomerase reverse transcriptase (mTert) expression marks slowly cycling intestinal stem cells. Proc Natl Acad Sci U S A. 2011;108:179–84.

9. Sangiorgi E, Capecchi MR. Bmi1 is expressed in vivo in intestinal stem cells. Nat. Genet. 2008;40:915–20.

10. Takeda N, Jain R, LeBoeuf MR, Wang Q, Lu MM, Epstein JA. Interconversion between intestinal stem cell populations in distinct niches. Science. 2011;334:1420–4.

11. Ayyaz A, Kumar S, Sangiorgi B, Ghoshal B, Gosio J, Ouladan S, et al. Single-cell transcriptomes of the regenerating intestine reveal a revival stem cell. Nature. 2019;569:121–125.

12. Bujko A, Atlasy N, Landsverk OJ, Richter L, Yaqub S, Horneland R, et al. Transcriptional and functional profiling defines human small intestinal macrophage subsets. Journal of Experimental Medicine. 2018;215:441–458.

13. Delfini M, Stakenborg N, Viola MF, Boeckxstaens G. Macrophages in the gut: Masters in multitasking. Immunity. 2022;55:1530–1548.

14. Bain CC, Bravo-Blas A, Scott CL, Perdiguero EG, Geissmann F, Henri S, et al. Constant replenishment from circulating monocytes maintains the macrophage pool in the intestine of adult mice. Nat Immunol. 2014;15:929–937.

15. Fawkner-Corbett D, Antanaviciute A, Parikh K, Jagielowicz M, Gerós AS, Gupta T, et al. Spatiotemporal analysis of human intestinal development at single-cell resolution. Cell. 2021;184:810–826. e23.

16. Burclaff J, Bliton RJ, Breau KA, Ok MT, Gomez-Martinez I, Ranek JS, et al. A proximal-to-distal survey of healthy adult human small intestine and colon epithelium by single-cell transcriptomics. Cellular and Molecular Gastroenterology and Hepatology. 2022;13:1554–1589.

17. Yan KS, Gevaert O, Zheng GXY, Anchang B, Probert CS, Larkin KA, et al. Intestinal Enteroendocrine Lineage Cells Possess Homeostatic and Injury-Inducible Stem Cell Activity. Cell Stem Cell. 2017;21:78–90 e6.

18. Herring CA, Banerjee A, McKinley ET, Simmons AJ, Ping J, Roland JT, et al. Unsupervised trajectory analysis of single-cell RNA-seq and imaging data reveals alternative tuft cell origins in the gut. Cell systems. 2018;6:37–51. e9.

19. Heylmann D, Rodel F, Kindler T, Kaina B. Radiation sensitivity of human and murine peripheral blood lymphocytes, stem and progenitor cells. Biochim Biophys Acta. 2014;1846:121–9.

20. Takeda N, Jain R, LeBoeuf MR, Wang Q, Lu MM, Epstein JA. Interconversion between intestinal stem cell populations in distinct niches. Science. 2011;334:1420–1424.

21. Montgomery RK, Carlone DL, Richmond CA, Farilla L, Kranendonk ME, Henderson DE, et al. Mouse telomerase reverse transcriptase (mTert) expression marks slowly cycling intestinal stem cells. Proceedings of the National Academy of Sciences. 2011;108:179–184.

22. Yan KS, Chia LA, Li X, Ootani A, Su J, Lee JY, et al. The intestinal stem cell markers Bmi1 and Lgr5 identify two functionally distinct populations. Proc Natl Acad Sci U S A. 2012;109:466–71.

23. La Manno G, Soldatov R, Zeisel A, Braun E, Hochgerner H, Petukhov V, et al. RNA velocity of single cells. Nature. 2018;560:494–498.

24. Aibar S, González-Blas CB, Moerman T, Imrichova H, Hulselmans G, Rambow F, et al. SCENIC: single-cell regulatory network inference and clustering. Nature methods. 2017;14:1083–1086.

25. Bettess MD, Dubois N, Murphy MJ, Dubey C, Roger C, Robine S, et al. c-Myc is required for the formation of intestinal crypts but dispensable for homeostasis of the adult intestinal epithelium. Mol Cell Biol. 2005;25:7868–78.

26. Kim MJ, Xia B, Suh HN, Lee SH, Jun S, Lien EM, et al. PAF-Myc-Controlled Cell Stemness Is Required for Intestinal Regeneration and Tumorigenesis. Dev Cell. 2018;44:582–596 e4.

27. Zimberlin CD, Lancini C, Sno R, Rosekrans SL, McLean CM, Vlaming H, et al. HDAC1 and HDAC2 collectively regulate intestinal stem cell homeostasis. FASEB J. 2015;29:2070–80.

28. Shiota M, Zoubeidi A, Kumano M, Beraldi E, Naito S, Nelson CC, et al. Clusterin is a critical downstream mediator of stress-induced YB-1 transactivation in prostate cancer. Mol Cancer Res. 2011;9:1755–66.

29. Xian L, Georgess D, Huso T, Cope L, Belton A, Chang YT, et al. HMGA1 amplifies Wnt signalling and expands the intestinal stem cell compartment and Paneth cell niche. Nat Commun. 2017;8:15008.

30. Oshima H, Kok SY, Nakayama M, Murakami K, Voon DC, Kimura T, et al. Stat3 is indispensable for damage-induced crypt regeneration but not for Wnt-driven intestinal tumorigenesis. FASEB J. 2019;33:1873–1886.

31. Sarper SE, Inubushi T, Kurosaka H, Ono Minagi H, Kuremoto KI, Sakai T, et al. Runx1-Stat3 signaling regulates the epithelial stem cells in continuously growing incisors. Sci Rep. 2018;8:10906.

32. Bain CC, Scott CL, Uronen-Hansson H, Gudjonsson S, Jansson O, Grip O, et al. Resident and pro-inflammatory macrophages in the colon represent alternative context-dependent fates of the same Ly6Chi monocyte precursors. Mucosal Immunol. 2013;6:498–510.

33. Lavin Y, Mortha A, Rahman A, Merad M. Regulation of macrophage development and function in peripheral tissues. Nat Rev Immunol. 2015;15:731–44.

34. Saeed S, Quintin J, Kerstens HH, Rao NA, Aghajanirefah A, Matarese F, et al. Epigenetic programming of monocyte-to-macrophage differentiation and trained innate immunity. Science. 2014;345:1251086.

35. Soucie EL, Weng Z, Geirsdottir L, Molawi K, Maurizio J, Fenouil R, et al. Lineage-specific enhancers activate self-renewal genes in macrophages and embryonic stem cells. Science. 2016;351:aad5510.

36. Francois A, Milliat F, Guipaud O, Benderitter M. Inflammation and immunity in radiation damage to the gut mucosa. Biomed Res Int. 2013;2013:123241.

37. Gupta VK, Jaskowiak NT, Beckett MA, Mauceri HJ, Grunstein J, Johnson RS, et al. Vascular endothelial growth factor enhances endothelial cell survival and tumor radioresistance. Cancer J. 2002;8:47–54.

38. Molla M, Panes J. Radiation-induced intestinal inflammation. World J Gastroenterol. 2007;13:3043–6.

39. Mintet E, Rannou E, Buard V, West G, Guipaud O, Tarlet G, et al. Identification of Endothelial-to-Mesenchymal Transition as a Potential Participant in Radiation Proctitis. Am J Pathol. 2015;185:2550–62.

40. Kiang JG, Olabisi AO. Radiation: a poly-traumatic hit leading to multi-organ injury. Cell Biosci. 2019;9:25.

41. Tian H, Biehs B, Warming S, Leong KG, Rangell L, Klein OD, et al. A reserve stem cell population in small intestine renders Lgr5-positive cells dispensable. Nature. 2011;478:255–9.

42. Hanzelmann S, Castelo R, Guinney J. GSVA: gene set variation analysis for microarray and RNA-seq data. BMC Bioinformatics. 2013;14:7.

43. Liberzon A, Birger C, Thorvaldsdottir H, Ghandi M, Mesirov JP, Tamayo P. The Molecular Signatures Database (MSigDB) hallmark gene set collection. Cell Syst. 2015;1:417–425.

44. Raudvere U, Kolberg L, Kuzmin I, Arak T, Adler P, Peterson H, et al. g:Profiler: a web server for functional enrichment analysis and conversions of gene lists (2019 update). Nucleic Acids Res. 2019;47:W191–W198.

45. Aibar S, Gonzalez-Blas CB, Moerman T, Huynh-Thu VA, Imrichova H, Hulselmans G, et al. SCENIC: single-cell regulatory network inference and clustering. Nat Methods. 2017;14:1083–1086.

46. Jin S, Guerrero-Juarez CF, Zhang L, Chang I, Ramos R, Kuan C-H, et al. Inference and analysis of cell-cell communication using CellChat. Nature communications. 2021;12:1088.

